# Single molecule sequencing of M13 virus genome without amplification

**DOI:** 10.1101/133843

**Authors:** Luyang Zhao, Liwei Deng, Gailing Li, Huan Jin, Jinsen Cai, Huan Shang, Yan Li, Haomin Wu, Weibin Xu, Lidong Zeng, Renli Zhang, Huan Zhao, Ping Wu, Zhiliang Zhou, Jiao Zheng, Pierre Ezanno, Qin Yan, Michael Deem, Jiankui He

## Abstract

Third generation sequencing is a direct measurement of DNA/RNA sequences at the single molecule level without amplification. In this study, we report sequencing of the genome of the M13 virus by a new single molecule sequencing platform. Our platform detects single molecule fluorescence by the total internal reflection microscope technique, with sequencing-by-synthesis chemistry. We sequenced the genome of M13 to a depth of 316x and 100% coverage. The consensus sequence accuracy is 100%. We demonstrated that single molecule sequencing has no significant GC bias.

## INTRODUCTION

Since the first two publications of the human genome sequences [1, 2], scientists around the world have embarked on a quest for next-generation sequencing (NGS) technologies. The resulting progress has revolutionized fields ranging from academic research to clinic diagnosis [3, 4]. Applications in the field of precision medicine (see review [5] for precise definition) include cancer diagnosis [6, 7] and inherited disease diagnosis [8, 9]. Progress in NGS technologies has brought, ethical questions as well [10, 11]. Promising applications include detection of pathogenic organisms [12, 13] and forensic sciences [14, 15]. At present, the cost of the sample library preparation process for NGS is still a significant part of the total cost of genome sequencing. Simple operation, cost effective sample preparation, generation of high throughput data, and more sensitive instruments are key requirements of the sequencing market in the future.

Since the early stages of DNA sequencing, single molecule (SM) sequencing is a key technological development. SM sequencing was first experimented in late 80s [16] and is now seen as the next step in the evolution of NGS [17]. Different SM sequencing technologies have rapidly developed over the past decade, with progress on read length, sequencing time, and data throughput. Principles of these technologies exhibit notable differences. Three companies and their own technologies are now well known: (i) the first true single molecule sequencing (tSMS) combining with sequencing-by-synthesis (SBS) [18] technology from Helicos Biosciences [19, 20] which is the technology we are improving; (ii) single molecule real time (SMRT) sequencing technology from Pacific Biosciences which provides super long read length (longer than 10k bases [21, 22]), but relatively low throughput; and (iii) Oxford Nanopore platform based on the direct electrical detection of single DNA molecule through α-hemolysin nanopores on which surface exonuclease enzyme molecules are attached [23], which provides long read (6k bases [21]) but limited accuracy and low throughput.

Despite the advantages of NGS platforms, the preparation of DNA libraries generally requires a preliminary step based on PCR amplification. This process introduces bias and ultimately can result in wrong interpretation of raw data sets [24, 25]. The Illumina sequencing platform, with which most current sequencing is performed, produces data sets showing uneven coverage and serious defects in GC-poor or GC-rich regions. Low coverage regions could be interpreted as sequencing errors by most current assemblers [26], and high coverage regions could be interpreted as repetitive sequences [27, 28]. Much effort has gone into improving protocols of library preparation to reduce or fully suppress GC bias [29, 30].

Advantages of the sample preparation for SM sequencing, combined with massively parallel short reads covalently captured onto an engineered surface, ideally fit the requirements of clinical diagnosis using DNA sequencing. Advantages of SM sequencing include (i) a simple and time-saving sample preparation consisting briefly of DNA shearing followed by poly-A tailing and 3′ end blocking steps, (ii) absence of base substitution introduced by the limited fidelity of DNA polymerases routinely used in PCR to amplify genomic DNA in samples, and (iii) the possibility of sequencing RNA molecules as well as DNA in order to investigate transcriptomic aspects of gene expression.

Our approach is devised to provide simple operation and high-throughput, unbiased data. Recently, we have demonstrated a direct targeted sequencing of cancer related gene mutations at the SM level [31]. In this paper, we describe the performance of our new GenoCare platform for SM sequencing without preliminary PCR amplification. The vector M13mp18 whose sequence is derived from the genome of the bacteriophage M13 was sequenced to the depth of 316x with 100% coverage. More importantly, no significant GC bias was observed.

## EXPERIMENTAL CONSIDERATIONS

### SAMPLE PREPARATION

**M13**: M13mp18 cloning vector was purchased from NEB, Beijing, China, and used as received. The sequence of the M13mp18 cloning vector is derived from the M13 phage [32] and contains 7249 bp. In this study, we used this cloning vector as DNA raw material to re-sequence, analyze, and compare with the reference sequence.

**Oligonucleotide Primers:** 5’ amine functionalized Poly-T oligonucleotides were purchased from Sangon and used as received.

M13 genomic DNA preparation process was illustrated in Figure 1.

**Figure 1.**
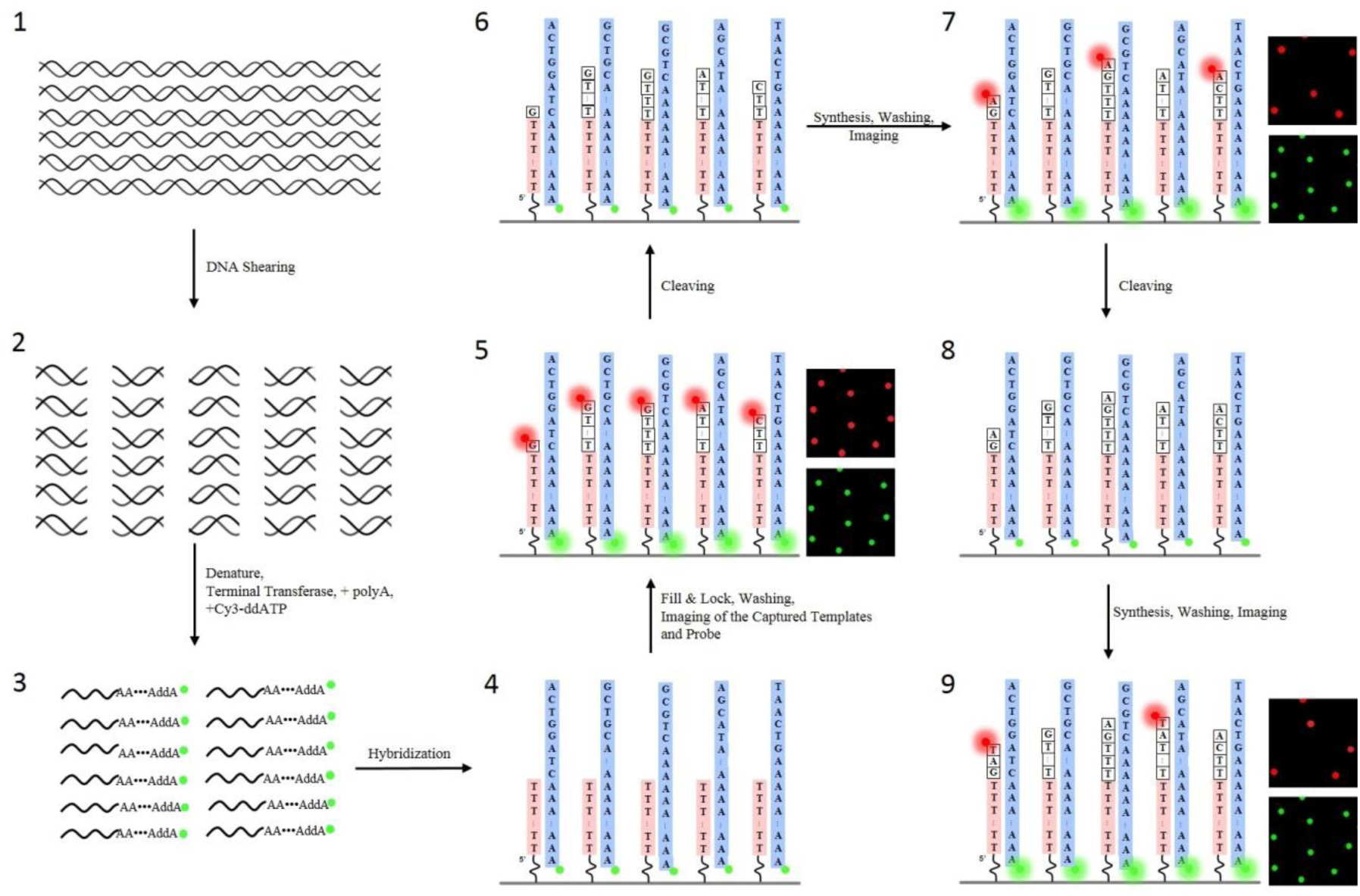
Sample preparation and sequencing process for single molecule sequencing of biological samples.

#### 1. DNA fragmentation

The M13mp18 cloning vector (from NEB, ref. N4018S) was used as raw DNA material to be sequenced by our platform. This cloning vector was first randomly fragmented into dsDNA fragments of about 200 bp using NEBNext® dsDNA Fragmentase® (from NEB, ref M0348S). Then, DNA fragments were purified using Agencourt AMPure XP beads (from Beckman, ref. A63881). The concentration of DNA was assessed by UV absorption using a Nanodrop 2000 device.

#### 2. Poly-A tailing and blocking

Multiple incorporations of 50-100 dATP at the 3′ end of ssDNA fragments from the cloning vector resulted in a poly-A tail. This reaction completed within 20 minutes. In a second step, poly-A tailed 3′ ends were blocked by incorporating the Cyanine 3 dideoxy ATP (Cy3-ddATP from PERKINELMER, ref. NEL586001EA). The blocking reaction completed within 30 minutes using the enzyme Terminal Transferase (from NEB, ref. M0315) such that the incorporation of reversible terminators at the 3′ end of the template strands was prevented.

## SURFACES AND TEMPLATE CAPTURE

### Surface Chemistry

Sequencing surfaces were prepared on 110×74 mm epoxy-coated glass coverslips (SCHOTT, Jena, Germany). Poly-T oligonucleotides were covalently bond to surface.

### Flow Cells

The above functionalized glass coverslip was assembled with a 1.0 mm thick glass slide by a pressure sensitive adhesive to form a flow cell. The flow cell has 16 channels, determined by the adhesive shape. For the M13 sequencing in this experiment, ∼0.5% of one channel was imaged.

### Template Capture (Hybridization)

The surface of the flow-cell was chemically modified by anchoring poly-T ssDNA strands at their 5′ end, in order to capture poly-A tailed strands from the library once they were injected inside the flow-cell at 55 °C. Then non-hybridized templates were washed away by 150 mM HEPES, 1X SSC and 0.1% SDS, followed by 150 mM HEPES and 150 mM NaCl.

## SEQUENCING REACTIONS

### The GenoCare Platform

All the sequencing reactions were implemented on the GenoCare platform The GenoCare is an automated single molecule sequencer with three major components: fluorescence imaging system, microfluidic system, and the stage to control the movement of sample. The imaging system is based on total internal reflection fluorescence (TIRF) microscopy [31].

### Fill & Lock

Since the hybridization of poly-T primer with poly-A tailed template may not be perfect, a step to fill the remaining dATP on the template with dTTP before the real sequencing process starts is necessary. After hybridization, the temperature of the flow-cell was lowered to 37 °C. The unpaired adenine nucleotides of poly-A tailed template strand were paired by multiple incorporations of natural thymine nucleotides at the 3′ end of primer strands. A mixture of dATP, dCTP, and dGTP reversible terminators were added to block further incorporation so that the template was locked in place and ready for sequencing.

### Nucleotide Addition

Reversible terminators were adopted in the sequencing-by-synthesis approach. They are modified nucleotides, which are composed of nucleotide triphosphates, a fluorophore (Atto647N), disulfide linker, and an inhibitor group. The design of the inhibitor effectively blocks the incorporation of next nucleotide before the cleavage of the disulfide bond of the previous reversible terminator.

The DNA extension was carried out at 37°C in Tris buffer containing polymerase, one of the four nucleotides and other salts. The components of this system are available with the use instructions from Direct Genomics.

## RESULTS AND DISCUSSION

### Sequencing Process

Our sequencing-by-synthesis (SBS) scheme is shown in Figure 1. Sample preparation is simple and fast, especially without amplification. M13 genomic DNA was sheared into fragments of ∼200bp, a length of 50∼100nt poly-A was added to the end, and blocked by ddATP-Cy3. Sequencing surfaces were chemically modified and covalently bound with poly-T, which can be hybridized with target DNA. Once annealed, residual dATP were filled with natural nucleotides, and locked with one reversible terminator. The sequencing of target DNA was then ready to commence.

The single molecule SBS process has been described elsewhere [31] Each cycle includes terminator incorporation, imaging, cleavage of fluorophore, and capping of residual bonds. The GenoCare platform adopts the total internal fluorescence microscopy (TIRF) for the observation of single molecules. The integration time was 200 ms to guarantee a good signal-to-noise ratio and reduce the photobleaching of dyes. We used 0.5% of one flow-cell channel to resequence the M13 virus genome as a demonstration of the GenoCare performance. We sequenced 80 cycles (40 quads of CTAG), and the images were analyzed (**Supporting Information**) to reach 100% coverage and an average depth of 316x for each base. The instrument run time was 9 hours, and sample preparation took 3 hours.

### Genome Coverage

About 100,000 reads were uniquely aligned to the reference genome, which accounted for 25.4% of the total reads. Reads were filtered by several criteria: 1) reads that were less than 13 bases after alignment were discarded, 2) reads that included a sequence exactly matching the terminator addition order and indicative of non-specific adsorption were also discarded, and 3) reads that could be mapped to multiple locations on the reference genome were excluded. From these data, we can determine the error rate. As is shown in Table 1, the dominant error was deletion (1.65%), followed by insertion (0.78%) and substitution (0.69%). Most of the reads aligned perfectly to the reference without any error, and the most error allowed per sequence by our algorithm was 3 (**Figure S1**). The average coverage depth for each base is 316x, and the minimum coverage is 14x (Figure 2a). When the average coverage depth reaches 10x, genome coverage rate climbs to 100% (Figure 2b). The Integrative Genomics Viewer (IGV) gives a clear picture of mapping against the known M13 genome reference (**Figure S2**).

**Figure 2.**
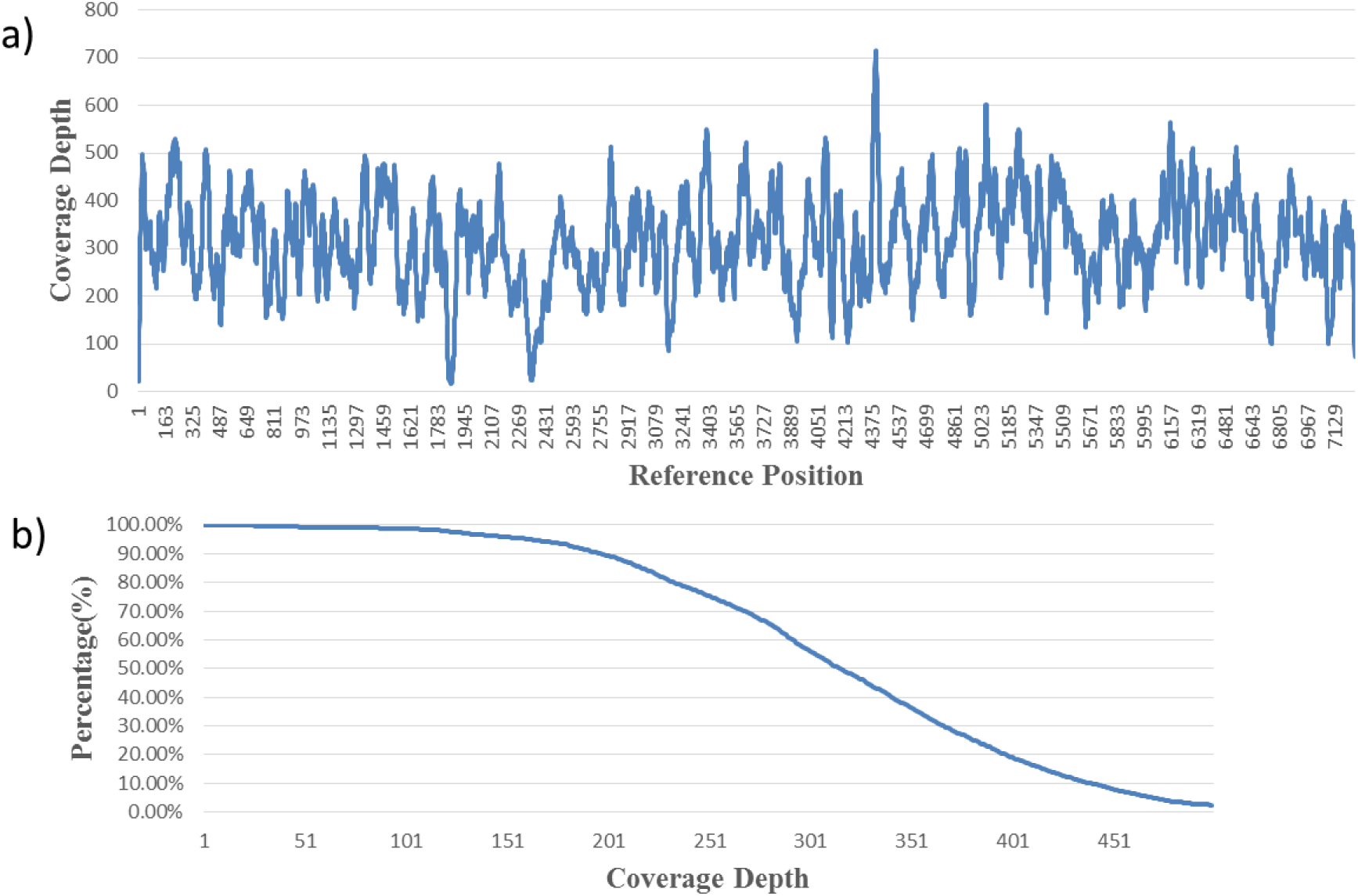
(a) Coverage depth for each base on M13 reference. The average coverage depth is 316x±96x. (b) Coverage rate as a function of coverage depth. 100% coverage was achieved when average coverage depth reached 10X.

**Table 1.** M13 genome sequencing statistics

| Cycles Sequenced | Average read length | Coverage |  |  | Total Reads | Mapped Reads | Unique Mapped Reads | Unique Mapped Ratio | Sub. Rate | Del. Rate | Ins. Rate |
| --- | --- | --- | --- | --- | --- | --- | --- | --- | --- | --- | --- |
|  |  | Average | Max. | Min. |  |  |  |  |  |  |  |
| 80 | 22 bases | 316x | 717x | 18x | 409491 | 104802 | 103990 | 25.4% | 0.69% | 1.65% | 0.78% |

### Read Length

The read length for this M13 sequencing run is shown in Figure 3. With the sequences less than 13 bases discarded due to poor unique mapping rate, this average read length is 22 bases (Table 1), given that only 80 base incorporation cycles were conducted. For the 7.2 kb M13 genome, a read length of 22 bases gives more than adequate specificity for alignment. Before alignment, the length distribution shows a peak at around 25 bases, while many reads were lost due to errors and non-uniqueness of mapping during alignment, causing a decrease of throughput and lowering of the average read length.

**Figure 3.**
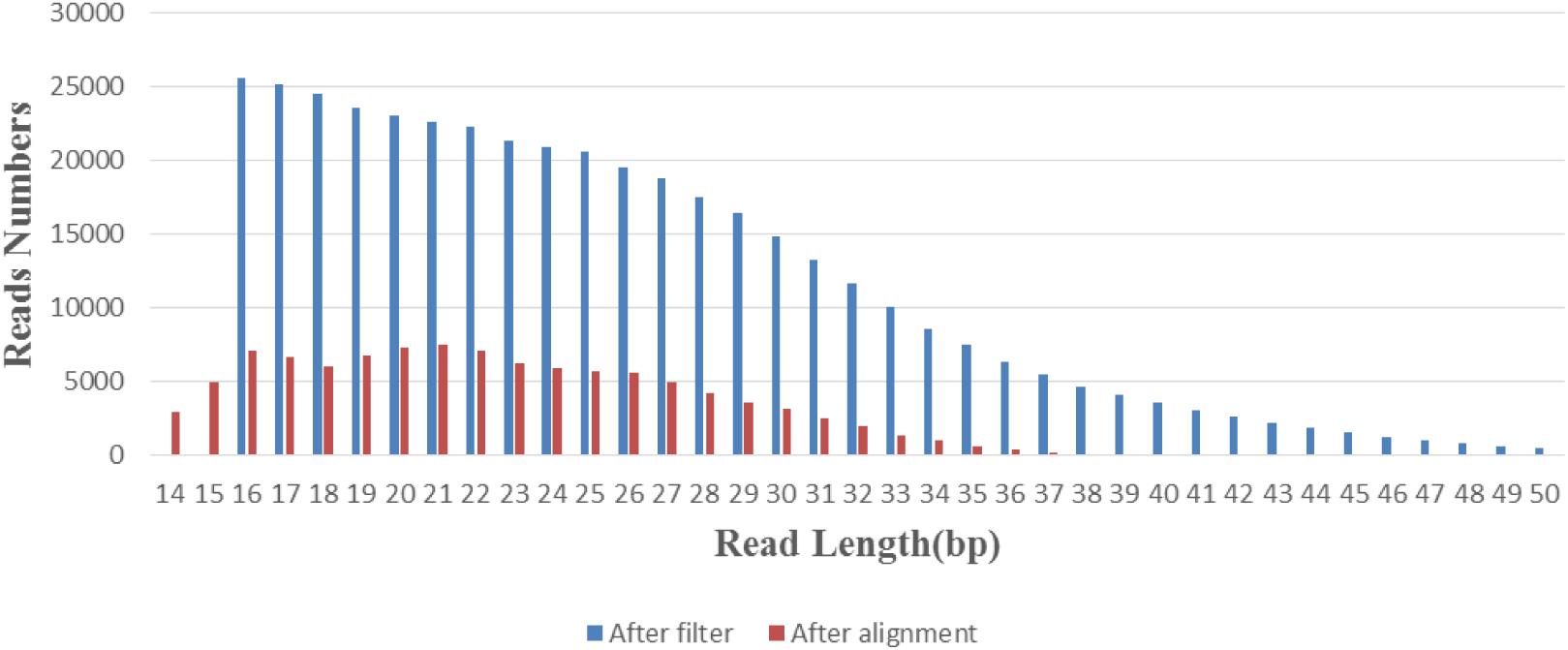
Read length distribution after length and repeat filters (blue bars) and after alignment (red bars).

### GC Bias

In accordance with the predicted effect of a PCR-free sample preparation, no obvious GC bias was observed under a window of 100 bases in which the GC content fluctuates in the range 22-69% (Figure 4a). The y-axis is an average of coverage depth in all 100-base windows with the same GC percentage. The distribution of base frequency in the reference as function of the CG content shows an almost identical shape to the depth distribution calculated from the sequencing result (Figure 4b and **Supporting Information**) and the R^2^ (goodness of fit) of those two curves is 0.9946, which indicates that no coverage bias is observed in this experiment.

**Figure 4.**
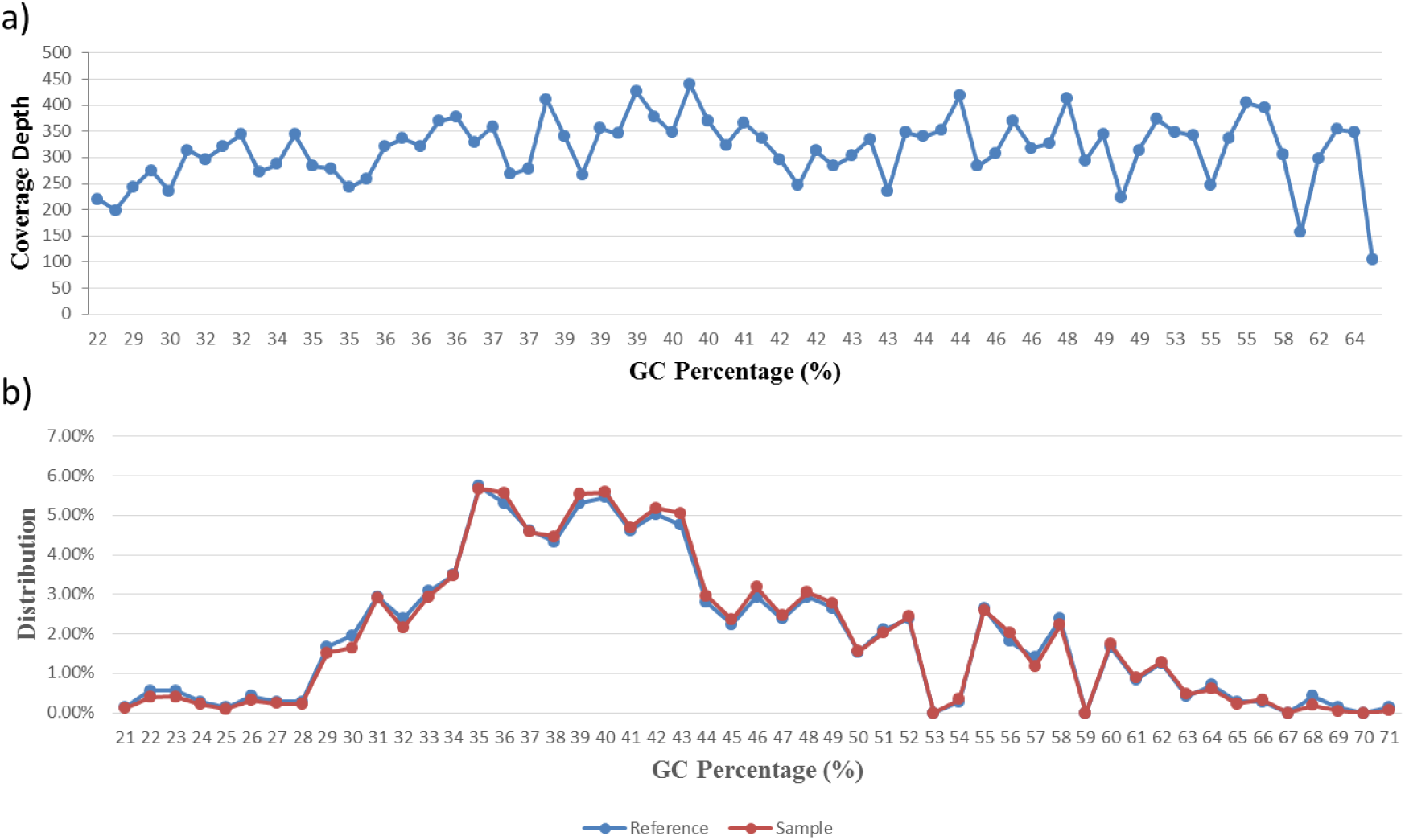
(a) Average depth distribution of all 100-base windows as a function of GC content. From GC content 22% to 69%, the average depth of each window on genome fluctuates in a small range. (b) GC patterns of reference genome (blue curve) and aligned reads (red curve).

## CONCLUSION

In this paper, we demonstrated the new GenoCare platform for single molecule sequencing. GenoCare is an automated desktop type of sequencer for dedicated use in the clinic. Compared to traditional next generation sequencing, sample preparation of single molecule sequencing is simpler and faster. Most importantly it does not require the use of PCR amplification, which effectively limits the GC bias. For example, on the Illumina system, a major NGS platform, it has been reported that GC bias leads to an uneven coverage or even no coverage of reads across the genome.

The cloning vector M13mp18 was sequenced on this new platform. A total of 80 cycles was run. Overall sequencing took 12 hours including sample preparation, instrument run time and data analysis. Eventually an average of 316x coverage depth and 22 bases read length were achieved. The consensus accuracy reached 100% once each base was sequenced at least 10 times. There was no apparent GC bias observed in this experiment, demonstrating the advantage of single molecule sequencing.

